# Improved blind demixing methods for recovering dense neuronal morphology from barcode imaging data

**DOI:** 10.1101/2021.08.10.455873

**Authors:** Shuonan Chen, Jackson Loper, Pengcheng Zhou, Liam Paninski

## Abstract

Cellular barcoding methods offer the exciting possibility of ‘infinite-pseudocolor’ anatomical reconstruction — i.e., assigning each neuron its own random unique barcoded ‘pseudocolor,’ and then using these pseudocolors to trace the microanatomy of each neuron. Here we use simulations, based on densely-reconstructed electron microscopy microanatomy, with signal structure matched to real barcoding data, to quantify the feasibility of this procedure. We develop a new blind demixing approach to recover the barcodes that label each neuron. We also develop a neural network which uses these barcodes to reconstruct the neuronal morphology from the observed fluorescence imaging data, ‘connecting the dots’ between discontiguous amplicon signals. We find that accurate recovery should be feasible, provided that the barcode signal density is sufficiently high. This study suggests the possibility of mapping the morphology and projection pattern of many individual neurons simultaneously, at high resolution and at large scale, via conventional light microscopy.

## 1 Introduction

Neuroscientists have long dreamed of obtaining simultaneous maps of the morphology of every neuron in a mammalian brain [1,2]. The ability to perform this very high-throughput neuron-tracing would enable better understanding of brain development, neural circuit structure, and the diversity of morphologically-defined cell types [3, 4]. In order to map the morphology at the scale of a mammalian brain, an ideal experiment would trace a large number of neurons simultaneously over a large brain region with high imaging resolution.

Current experimental methods for neuronal tracing can be placed along a spectrum spanning two disparate spatial scales. On the one hand, techniques based on Electron Microscopy (EM) obtain nanometer-level resolution within small regions. On the other hand, light microscopy and molecular techniques can map morphology over whole-brain scales, albeit with either low spatial resolution or limitations on the number of neurons that can be mapped simultaneously. Figure 1 sketches the current landscape, and highlights a gap in the state of the art: mapping many neurons simultaneously, with high resolution, over large fields of view (or potentially even the whole brain).

**Figure 1:**
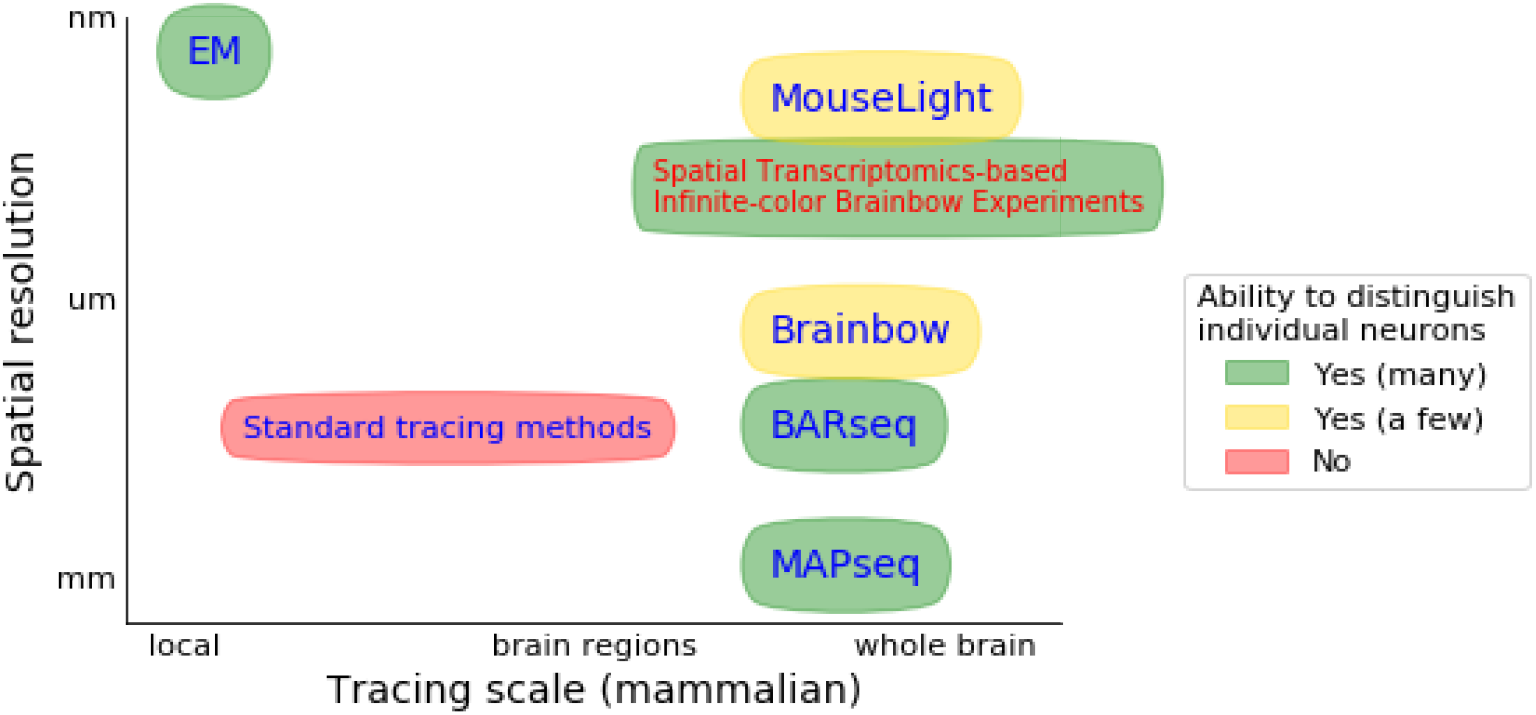
Some existing methods for neuronal morphology mapping. Here we non-exhaustively summarize the landscape of existing neuronal tracing methods. The horizontal axis indicates the typical scale of these neuronal tracing methods, ranging from local tracing to mapping axonal projections across the whole brain; the vertical axis indicates optical resolution in the typical images obtained from these methods. The ellipse color represents the number of neurons that can be traced and distinguished; red indicates that the method cannot readily distinguish between spatially nearby neurons, yellow indicates that the method can be used to trace several individual neurons, and green indicates the possibility of simultaneously mapping thousands or millions of neurons. Here ‘standard tracing methods’ refer to approaches such as Golgi staining and viral tracers; cf. Section 2 for more details. Note that many of these methods rely on conventional optical microscopes, and their resolution can potentially be increased via expansion microscopy and/or super resolution microscopy, though this incurs additional experimental costs. In this paper we focus on the possibility of extending molecular barcoding methods by increasing the signal density to allow for more fine-grained morphological reconstructions; we call such experiments Spatial Transcriptomics-based Infinite-color Brainbow Experiments (STIBEs).

Molecular barcoding methods may be able to fill the gap [5,6] with ‘infinite-color Brainbow’ experiments: each cell is labeled with a unique random barcode which can be visualized through conventional light microscopy. If we think of each barcode as providing a unique ‘pseudocolor’ for each cell, then conceptually this method ‘colors’ the brain with a potentially unbounded number of random colors, generalizing the original Brainbow approach [7], which can trace long-range neuronal morphology but was limited to a relatively small number of random colors (though see e.g. [8] for more recent developments).

Here we use simulations to investigate a class of methods we dub Spatial Transcriptomics-based Infinite-color Brainbow Experiments (STIBEs); can such experiments obtain precise morphological reconstruction and long-range tracing for many neurons simultaneously? This class of experiments uses techniques from the spatial transcriptomics community to label cells with unique fluorescent barcodes, in-situ; the BAR-seq method is a representative example [5]. STIBEs work by creating a collection of transcripts containing barcode sequences; these transcripts are amplified into target molecules known as ‘amplicons.’ Amplicons are generally less than one micron in diameter, and the barcode associated with each amplicon can be identified over a series of rounds of imaging. In an ideal STIBE, all amplicons within a given cell carry the same barcode, no two cells carry amplicons with the same barcode, and amplicons are only present within cells. By identifying collections of amplicons which carry the same barcode, one can thus trace many neurons across the entire brain.

Previous experiments have already demonstrated that STIBEs can be used to recover the locations of somas and projection sites of many cells (at lower spatial resolution) [5, 9]. In contrast, here we focus on STIBEs that could achieve micron-resolution morphological reconstruction of many cells simultaneously, potentially across the whole brain. This would be a valuable advance for several reasons. First, cell morphology measured at this scale provides a great deal of information about cell type in different layers and regions of the brain [10, 11]. Second, although micron-level resolution does not permit exact connectome reconstruction, it does yield a matrix of *potential connections* between neurons (i.e., which cells might be connected to each other). Such matrices would be invaluable for downstream applications such as functional connectivity mapping (by providing constraints on possible connections) [12] or cell type inference [13–15].

To accurately simulate the imagestacks that would be generated by STIBEs, we use two sources of existing data. First, we use densely-reconstructed EM data to give us the shapes and spatial relationships between axons and dendrites in a small cortical volume. Second, we replicate the noise and signal structure we might expect, based on results from current cellular barcoding experiments [5, 6]. We investigate how varying parameters of STIBEs affects the accuracy of neuronal tracing; for example, we consider various imaging resolutions and various densities of the amplicons present within cells. We find that selecting an appropriate density of amplicons is a balancing act. We cannot recover precise morphological reconstructions without sufficient density, but very high density can lead to incorrect amplicon identification along thin processes (especially in the presence of relatively low optical resolution).

Our main technical contribution here is to develop improved demixing algorithms that enable improved barcode recovery in the high-density regime, leading to improved morphological recovery. These algorithms are based on the BarDensr model of spatial transcriptomics data [16] (see also [17], as well as [18,19]), which was originally developed to detect barcode signals from spatial transcriptomics images when the barcode library is known. We have extended this method to the more challenging ‘blind demixing’ setting, so that it is applicable to STIBEs where the barcode library is unknown or intractably large. We further developed a Convolutional Neural Network (CNN) model to reconstruct morphology for each demixed barcode, ‘connecting the dots’ between the discontiguous fluorescent signals produced by amplicons.

In what follows, we contextualize STIBEs among the family of other neuronal tracing methods, describe our methods for simulation and image analysis, and finally report our findings on the feasibility of STIBEs for the morphological reconstruction of neurons.

## 2 Related work

The history of neuronal tracing techniques is long and complex [20]; here we give a brief overview of some techniques most directly relevant to our approach. Figure 1 presents a visual overview.

At one extreme of the precision-scale spectrum, Electron Microscopy (EM) is the state-of-the-art tool for visualizing a small region of brain with very high precision [14, 15,21–25]. However, it is not yet suitable for dense mapping of long-range neuronal projections: the imaging (and computational segmentation) process for these experiments is challenging, and it is not yet feasible to use EM to densely trace many neurons across the mammalian brain. It may be possible to refine EM techniques to overcome these challenges, but this is not our focus in this paper (cf. [2] for further discussion).

At the opposite extreme of the precision-scale spectrum, we have standard tracing approaches utilizing light microscopy. Despite significant effort [26], it remains infeasible to segment and trace thousands of densely-packed individual neurons visualized with a single tracing marker. The ‘MouseLight’ project [27] is an exemplar of this ‘classical’ approach. This is a recently developed platform for tracing the axonal arbor structure of individual neurons, built upon two-photon microscopy and viral labeling of a sparse set of neurons, allowing long-range neuronal tracing. Unfortunately, the number of neurons that can be traced at once is currently limited to a hundred per brain sample [28] (though more advanced microscopy technology, such as super-resolution microscopy [29–33], and/or Expansion Microscopy [34, 35], could potentially be deployed to simultaneously trace more individual neurons with higher resolution).

‘Brainbow’ methods [7,36] introduce random combinations of fluorescent markers to facilitate tracing of multiple cells. The number of neurons that can be uniquely labeled and identified by the original method was limited to several hundreds, because of limitations on the number of distinct colors that can be generated and distinguished. Recent studies have focused on reducing these bottlenecks [8, 37, 38].

More recently, molecular barcoding methods have been developed to effectively remove this bottleneck on the number of distinct unique ‘color’ labels that can be assigned and read from different cells. Multiplexed Analysis of Projections by Sequencing (MAPseq) is one early example of this approach, developed to study long-range axonal projection patterns [9]; however, the spatial precision of this approach was not sufficient to capture neuronal morphology. More recently, BARseq [5, 6] combines MAPseq and *in situ* sequencing technology to obtain higher spatial resolution. Instead of micro-dissecting the brain tissue, this method uses fluorescent microscopy to detect the barcode signal from the intact tissue, similar to recent spatial transcriptomics methods [39, 40]. In theory, BARseq should be capable of satisfying all the desirable features of neuronal tracing: uniquely labelling large numbers of individual neurons, representing them with high spatial precision, and tracing them across long distances through the brain. Spatial transcriptomics technology is advancing quickly; just this year [41] achieved single-cell-resolution transcriptomic assays of an entire embryo. As this technology advances, techniques like BARseq can only become more effective. However, to date, this technology has only been used with a relatively small number of amplicons per cell, prohibiting accurate morphological reconstruction. In theory, there is nothing preventing new experiments with higher density, to achieve higher-resolution morphology mapping; this paper will explore this idea systematically.

Finally, we note the related paper [42]; this previous simulation work focused on a different experimental context, specifically expansion microscopy and cell membrane staining with fluorescent markers. In the current work we are interested in exploring the frontier of morphology mapping using only molecular barcoding images and conventional light microscopy. Introducing expansion microscopy and additional fluorescent labels could potentially improve the resolution of the methods studied here, at the expense of additional experimental complexity.

## 3 Methods summary

We here summarize the three main procedures used in this work; full details can be found in Appendix A.

### Data simulation

In order to investigate feasible experimental conditions for densely reconstructing neuronal morphology, we simulated a variety of STIBEs. The simulation process is illustrated in Figure 2. We started with densely segmented EM data [14,15,43]. Next we generated a random barcode library, assigning random barcodes to each cell in the field of view (FOV). We simulated amplicon locations according to a homogeneous Poisson process within the voxelized support of each cell. The final imagestack was generated by assigning a colored spot to each barcode location in each imaging frame, and then pushing the resulting high-resolution 3D ‘clean’ data through an imaging model that includes blurring with a point spread function, sampling with a lower-resolution voxelization, and adding imaging noise, to obtain the simulated observed imagestack.

**Figure 2:**
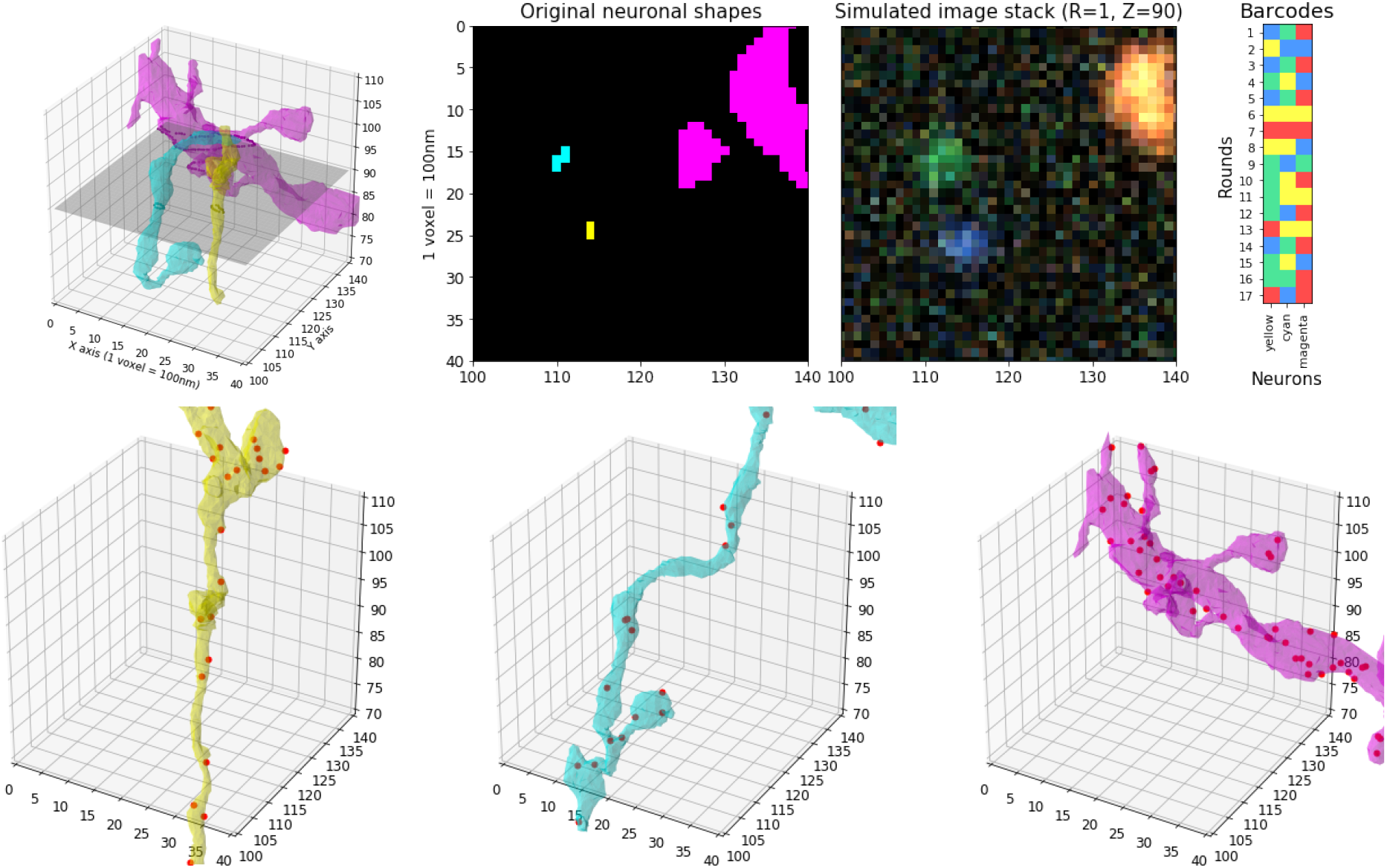
Simulation process illustration. **Top left:** Three neuronal segments in a small region (40 × 40 × 40 voxels with 100 × 100 × 100 *nm* voxel size). The original EM data is much more densely packed than what is shown here; for illustration purposes, we only show three neurons among many. **Top middle:** A 2D slice with each neuronal segment uniquely colored (left) and the simulated imagestack at the same plane for the first sequencing round (right). The imagestack colors do not correspond to neural segment colors, but rather to the corresponding fluorescent barcodes (top right). A video corresponding to this plot, showing multiple z-planes, can be found at this link. **Bottom:** Amplicons are uniformly simulated within neuronal segment volumes; simulated amplicons are shown as red dots.

### Barcode estimation

To use a STIBE for morphological reconstruction, one must estimate the barcodes present in a field of view (each barcode corresponding to one cell) and the locations of each amplicon corresponding to each barcode (these amplicon locations pointilistically trace the morphology of each cell). The barcode library used to infect the neurons is unknown in such data. In theory, one could sequence the viral library, but this may yield a set of barcodes which is too large to be useful. Indeed, the number of barcodes present in a field of view is much larger than the number of barcodes which actually infect cells [44]. For the purpose of morphology reconstruction – where only a small number of cellular barcodes exist in a region of interest – we must develop an approach to ‘learn’ the local barcode library from the images themselves. We found that there are effectively two distinct regimes for barcode and amplicon recovery. In the ‘sparse’ regime, where imaging resolution is high and the amplicon density is low, the signal in most imaging voxels is dominated by at most one amplicon. Thus we can estimate the barcode library simply by searching for bright, ‘clean’ imaging voxels displaying a single amplicon signal to estimate the amplicon locations using the resulting barcode library. In the ‘dense’ regime (with high amplicon density and/or low imaging resolution) it is harder to find voxels dominated by a single amplicon, and the simple approach described above breaks down. Instead, we have found that an iterative constrained non-negative matrix factorization approach is more effective: given an initial (incomplete) estimate of the barcode library, we estimate the corresponding amplicon locations, then subtract away the estimated signal corresponding to these amplicons. This sparsens the remaining image, making it easier to detect more barcodes that might have been obscured in previous iterations. After augmenting the barcode library with these previously undetected barcodes, we can re-estimate the amplicon locations and iterate between these two steps (updating the barcode library and estimating amplicon locations in alternating fashion) until a convergence criterion is satisfied.

### Amplicon estimation and morphological reconstruction

Given an estimated barcode library, our primary goal is to reconstruct the morphology of each neuron labeled by a barcode in this library. Our starting point in this endeavor is the ‘evidence tensor’ (see Appendix for a precise definition), which summarizes our confidence about the presence of each barcode at each voxel location. This evidence tensor is used as the input to two algorithms: alphashape [45] and Convolutional Neural Networks (CNNs), which are trained to estimate the shape of each neuron from the evidence tensor.

## 4 Results

Below we report our ability to recover barcodes and reconstruct morphology in two different simulation regimes: the ‘high-resolution, dense-expression’ regime, in which every neuron is labeled and imaged with sub-micron resolution; and the ‘low-resolution, sparse-expression’ regime, in which only about 1% of neurons are labeled and imaged with micron-resolution imaging. The first regime (‘high-resolution simulation’ for short) represents an ideal setting. Experiments in the second regime (‘low-resolution simulation’ for short) would be considerably cheaper to perform, with lower optical resolution facilitating faster image acquisition over large FOVs; however, the resolution in this simulation is low enough that many cellular barcodes may occupy a single voxel. To enable accurate recovery in this regime we must also assume that the cell capture efficiency is relatively low: i.e., only a fraction of neurons express a barcode.

### 4.1 High-resolution, dense-expression simulations

In our first set of simulations, we assume diffraction-limited imaging with an isotropic point spread function (PSF). We further assume that the viral infection has a perfect efficiency; in this extreme case, all the neurons captured by EM in this brain region are labeled with a unique barcode. The simulation process is illustrated in Figure 2. With these assumptions fixed, we investigated a range of experimental parameters, varying the amplicon density as a key parameter of interest. We denote this density by λ and measure it with units of amplicons per cubic micron of neuronal volume. We investigated densities from 1 amplicon per cubic micron (i.e., λ = 1) to 200 amplicons per cubic micron (i.e., λ = 200). For each density variant we investigated our ability to recover barcodes. The simulation results using three of these densities (1, 10 and 100 amplicons/*μm*^3^) are summarized in Figure 3. More details on the simulation process can be found in Appendix A.1.

**Figure 3:**
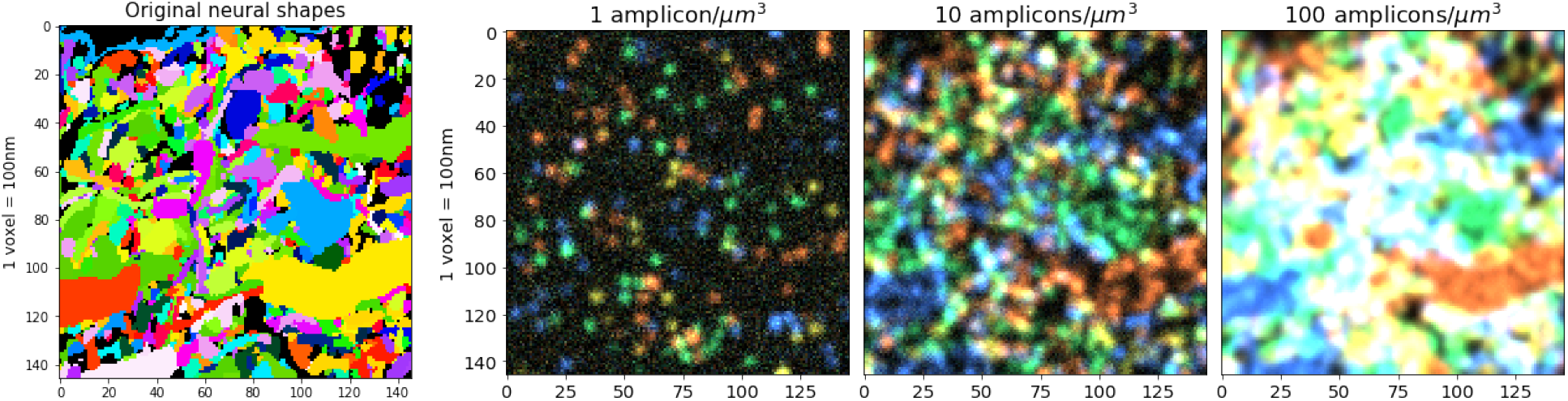
High-resolution, dense-expression simulation with varying amplicon densities. **Left:** 2D slice of the original neuronal shapes; each neuron is visualized with a different random color. **Right:** Corresponding slice of the simulated imagestack, considering several different density values. Note that the imagestack colors do not correspond to the neuronal visualization colors on the left; the imagestack colors arise from their corresponding barcodes. A video showing multiple z-planes can be found at this link.

#### 4.1.1 Barcode estimation

To estimate the barcode library present in a STIBE, we take advantage of a special structure in the barcodes of typical spatial transcriptomics experiments: each fluorescent barcode is designed so that exactly one channel is active in each imaging round. This suggests a simple approach for recovering the barcode library: search for voxels where one channel is much brighter than all others in every round. This approach is successful in some cases. Barcode discovery using this simple approach is easiest with a medium density of amplicons. In the low-density case, with λ =1 amplicon/*μm*^3^, we were able to find 847 out of 975 barcodes present in tissue using this naïve approach, representing a detection rate of 86.87%, with 0 false positives (however, as we discuss later, our ability to reconstruct neuronal morphology in this low-density case is quite limited). Increasing the density to 10 transcripts per cubic micron (λ = 10 amplicons/*μm*^3^), we are able to detect 945 barcodes (96.92% detection rate) with 2 false positives. Note that barcode recovery rates drop in the very low amplicon density regime, since some small cells in the FOV may not express enough amplicons for reliable detection.

As shown in Figure 4, even with medium density the signal can be significantly mixed; that is, a single voxel can display signal from multiple cells, even under the assumption of high-resolution imaging system. When this occurs, correctly identifying the barcodes becomes challenging even manually. Increasing the density further to 100 transcripts per cubic micron (λ = 100 amplicons/*μm*^3^), this problem becomes severe: the naïve method can detect only 604 barcodes, representing a detection rate of 61.95%, with zero false positives.

**Figure 4:**
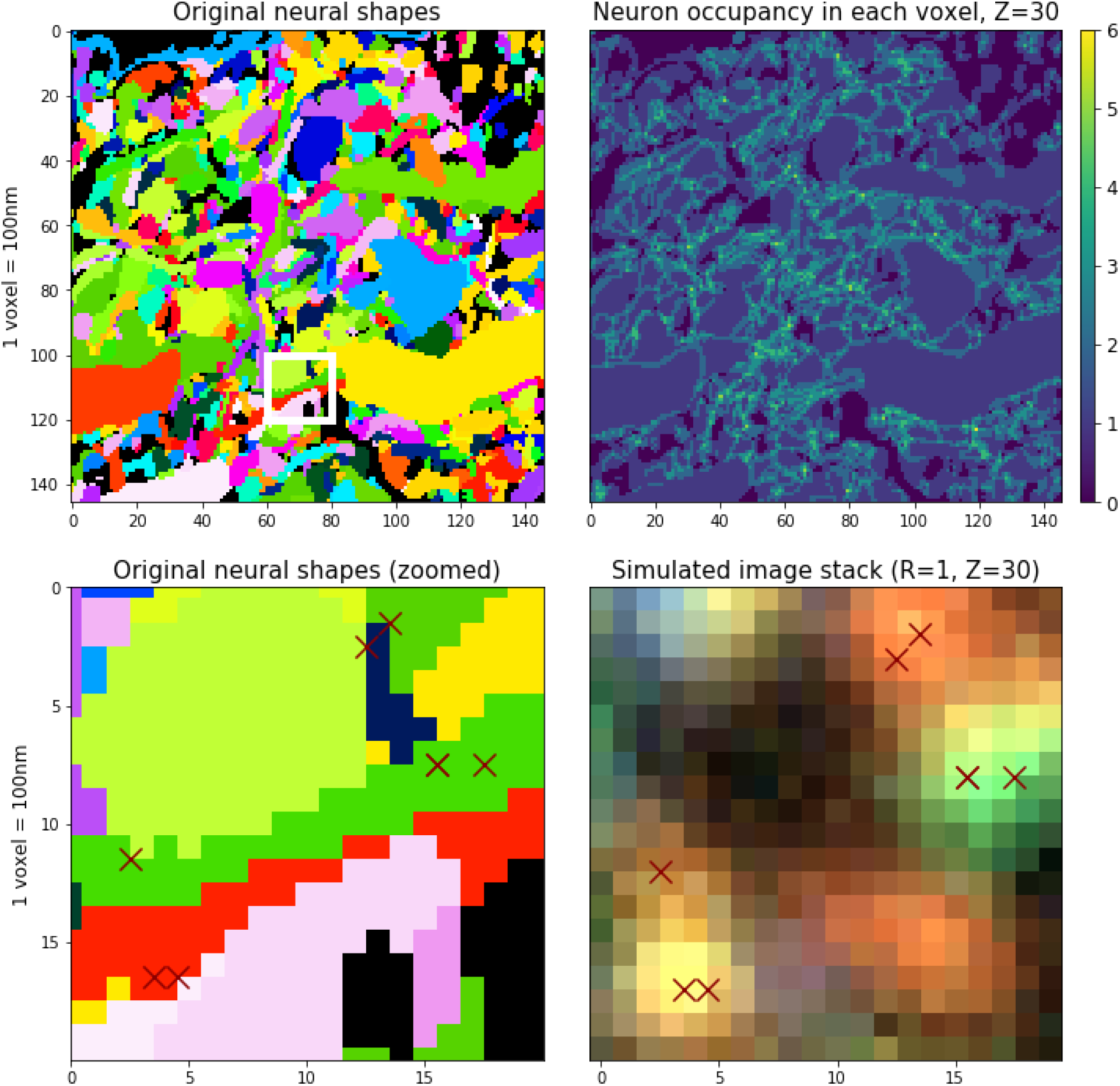
Limited optical resolution leads to signal mixing between neighboring neurons even in the high-resolution setting. **Top left:** Neuronal shapes in high-resolution, dense-expression data, as seen in Figure 3. **Top right:** The number of distinct neurons that contribute signal to each voxel. In some densely-packed regions, each voxel might mix signal from as many as six cells, due to optical blur and the voxelization process. **Bottom left:** Zoomed-in 20 × 20 region of the original neuronal shapes (the white rectangular region from the top left panel). **Bottom right:** Simulated imagestack in the same region, illustrating the challenge of uniquely assigning amplicons to neurons (λ = 10 amplicons/*μ*m^3^ here). Simulated amplicon locations within this z-plane are shown with red crosses; other amplicons centered outside of this z-plane contribute additional signal.

In short, we found the naïve barcode recovery method inadequate when the amplicon density is high, even under idealistic assumptions on optical resolution. In order to recover most of the barcodes, we need to demix the accumulated signal emitted by many barcodes contributing to a single voxel. We adopted a novel iterative approach based on non-negative matrix factorization, detailed in Section A.2.2 and Algorithm 3. The results of this iterative barcode discovery approach under different amplicon densities are shown in Figure 5. In all cases we chose thresholds to ensure at most 3 false positives. We found that 2-3 iterations consistently improve performance, yielding 30% more detections when the amplicon density is very high (λ = 100 amplicons/*μm*^3^).

**Figure 5:**
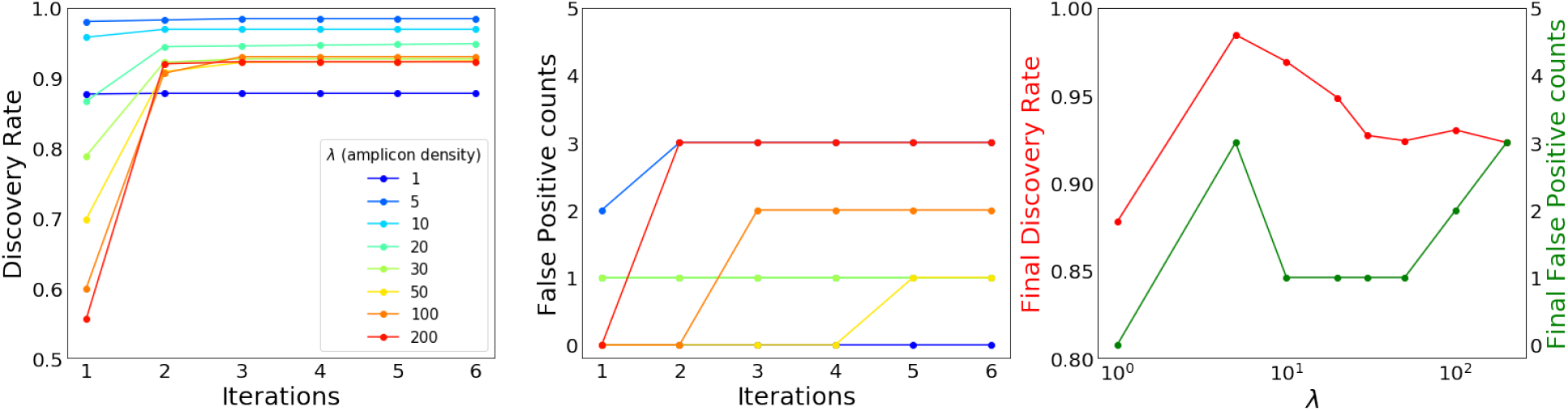
An iterative non-negative matrix factorization approach enables barcode discovery in the high-amplicon-density regime for high-resolution, dense-expression simulations. The left panel shows barcode discovery rate, as a fraction of the 975 neurons present in the simulations. The middle panel shows the False Positive counts. These two panels show these metrics evaluated on STIBEs with various densities of amplicons (λ, colored lines) analyzed using different numbers of iterations. We see that multiple iterations improve recovery accuracy when the density of amplicons is high. The right panel shows the final discovery rate (vertical axis on the left, red) as well as the final False Positive counts (vertical axis on the right, green) after six iterations, as a function of amplicon density (at log-scale, horizontal axis). Recovery accuracy peaks at λ values near 5 to 10 amplicons/*μ*m^3^, where amplicons are neither too dense (where too much optical overlap leads to decreases in recovery accuracy) nor too sparse (where there is simply not enough information to recover some barcode identities).

#### 4.1.2 Amplicon estimation and morphological reconstruction

In order to estimate the morphology corresponding to each recovered barcode, we start by estimating the density of each barcode at each voxel; we refer to this estimate as the ‘evidence tensor’ (cf. A.3). Note that this ‘evidence tensor’ is computed using barcodes recovered from the imagestack (rather than the ground-truth barcodes used for creating the simulations). Figure 6 compares the original neuronal shape and the corresponding evidence tensor; we see that the evidence tensor can roughly capture the shape of the original neuron.

**Figure 6:**
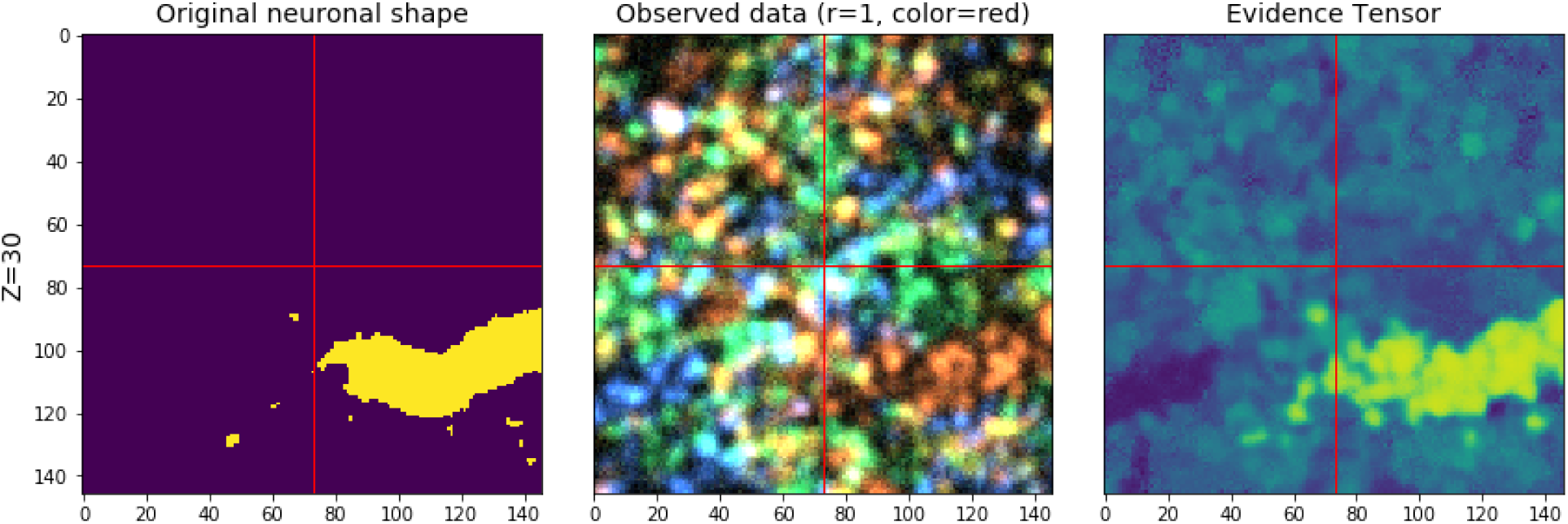
The evidence tensor is used to match barcodes to voxels. The ground-truth morphology at a selected z-plane for an example neuron is shown on the left; an imagestack from one round at the corresponding z-plane (λ = 10 amplicons/*μ*m^3^) is shown in the middle; on the right, the corresponding z-plane of the evidence tensor for the barcode corresponding to the neuron on the left is shown. Note that the evidence tensor is bright around the locations of the underlying neuron, as desired.

We also investigated neuronal tracing with low, medium and high-density variations in Figure 7. The evidence tensors computed from medium-density variation can capture the original morphology relatively well, by correctly identifying the amplicon locations (the third row). However, those learned from low-density variations cannot capture the original morphology as clearly (the second row). The rough path of the neuron can still be identified, but spines are generally missed from dendrites and sub-micron resolution is not achieved. The high-density variation results were similar to the medium-density variation, but with more continuous shape reconstruction (the bottom row). However, they tend to miss the detailed shapes, as well as a portion of the barcodes, due to overlaps of the simulated amplicons in such high-density images.

**Figure 7:**
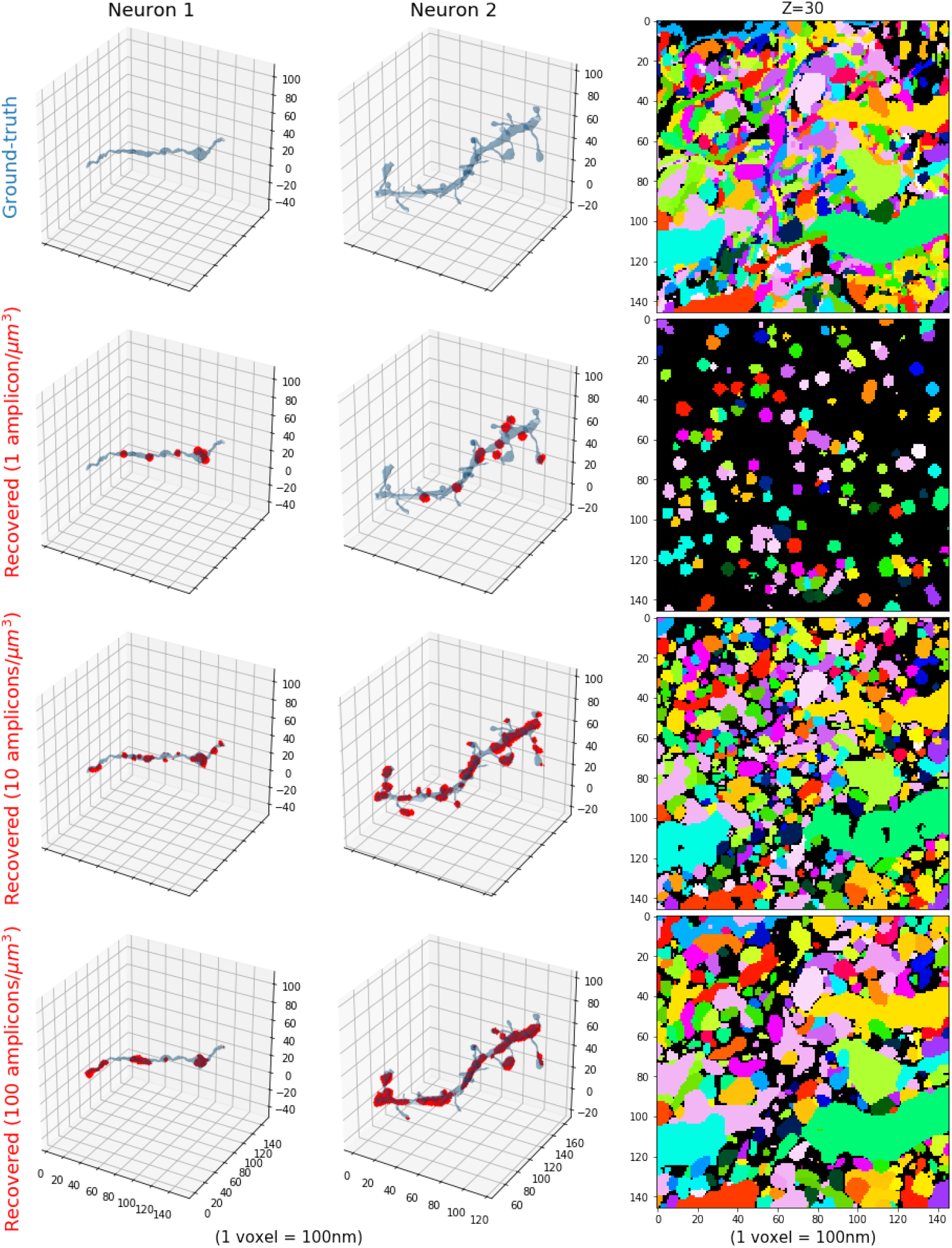
Neuronal morphology reconstruction from high-resolution, dense-expression simulations, with 1, 10 and 100 amplicons per cubic micron. Given an imagestack from a STIBE, how well can we recover the morphology of individual neurons? The first column visualizes one cell, the second visualizes another, both shown in blue shaded regions. The first row indicates ground-truth morphology of the cells. The second row shows our ability to locate points inside the cells using a high-resolution STIBE with λ = 1 amplicon/*μ*m^3^; identified locations are shown in red in the first two columns, and via unique colors in the third column. The final two rows visualize the same information for a STIBE with λ = 10 amplicons/*μ*m^3^ and λ = 100 amplicons/*μ*m^3^. **Right column**: On top, we visualize many cells in a single slice of tissue, each indicated with a unique color. The following rows show the recovered neuronal shapes based on the binarized evidence vector with threshold 0.7. Colors match the right top panel. A video showing multiple z-slices corresponding to the right panel can be found at this link.

Next we want to estimate the morphology of each neuron from the evidence tensor. We experimented with two approaches here. The first approach applies the alphashape algorithm [45] to a thresholded version of the evidence tensor; this enables better reconstruction of thin portions of the neuron by incorporating our knowledge that neurons occupy contiguous regions in space. Alphashape is a classical method to reconstruct morphology from a point cloud, obtained by computing Delaunay simplices and removing simplices with large circumradii. Figure 8 demonstrates that alphashape is able to estimate the continuous shape of neurons by ‘connecting the dots’ in the evidence tensor. However, it tends to over-estimate the shape, perhaps because alphashapes can only construct linear connections between pairs of detected signals. The second approach uses Convolutional Neural Networks (CNNs) applied to the evidence tensor. This method outperforms alphashape, as shown in Figure 8.

**Figure 8:**
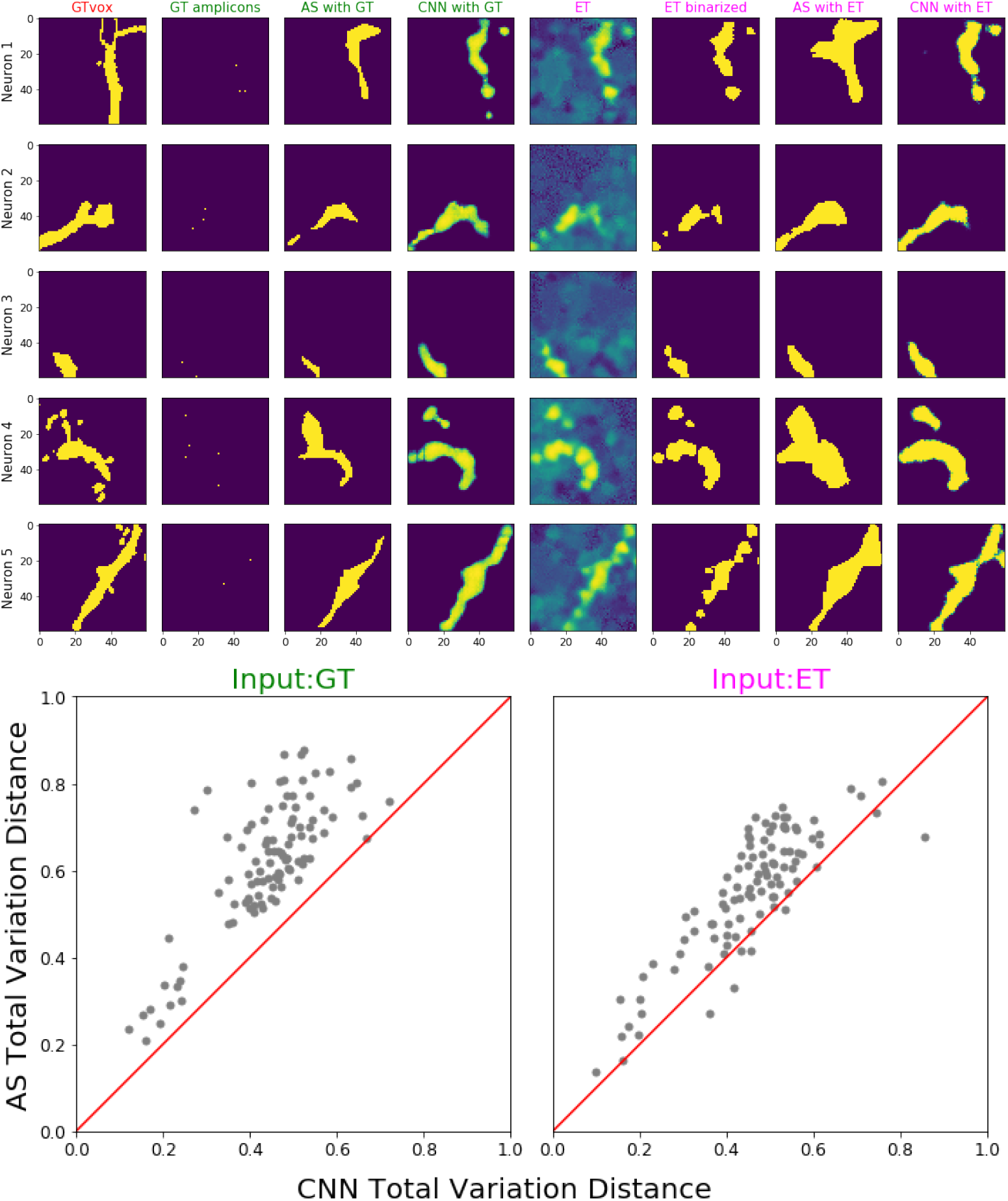
Comparing the morphology reconstruction for high-resolution, dense-expression simulations, using alphashape (AS) and CNN, with either ground-truth (GT) amplicons or evidence tensor (ET) as input. **Top:** Morphology prediction results for 5 examples. The first column shows the ground-truth morphology at one z-plane. The plane was chosen so that the voxel area occupied by each neuron is largest. All the following columns show the same z-plane as the first column. The next three columns show the prediction results using alphashape or CNN, when the ground-truth amplicons are used as input (indicated in green titles). The following four columns show the prediction results using alphashape or CNN, when the evidence tensors are used as input (indicated in magenta titles). Evidence tensors are first calculated using the correlation method described in Appendix A.3.1, and before input into alphashape, they are binarized with threshold 0.7, as shown. **Bottom:** Total variation distance (see Appendix A.3.2) on 100 testing examples, including the five shown on the top panel, comparing the results from alphashape prediction on the input (y-axis) and CNN prediction on the input (x-axis). Each dot represents one testing example. The red lines indicate where the two methods have the same performance. Lower values are better; the CNN significantly outperforms the alphashape reconstructions here. A video of the reconstruction result (across z-planes) can be found at this link.

In summary, morphological tracing is difficult with low-density STIBEs, but with sufficient density it is feasible to achieve sub-micron resolution. A thresholded version of the evidence tensor creates a satisfactory summary of the morphology, but it takes the form of a collection of discontiguous regions. To incorporate our knowledge that neurons occupy contiguous regions in space, we can use alphashape (which incorporates this knowledge directly) or CNNs (which learn from data); CNNs yield the best estimates.

### 4.2 Low-resolution, sparse-expression simulations

Next we investigate whether data from lower-resolution optical systems can still be used to reconstruct neuronal morphology. Here, we assume the optical system is less sensitive: the resolution is lower and the PSF is larger. We also assume the viral infection rate is lower, infecting only 1% of all the axonal segments and discarding all dendritic segments (emulating barcode injection at one site and axonal tracing at another site). This lower infection rate more closely mimics the low labeling efficiency by the current viral barcoding technologies. We assume amplicons can be produced with a density that is roughly consistent with the current experimental technologies, namely 0.08 amplicons per micron length [5]. More details on the low-resolution simulation can be found in Appendix A.1.

The left column of Figure 9 illustrates the difficulties in analyzing this kind of STIBE. The low optical resolution and high amplicon density often cause multiple amplicons belonging to different axons to occupy the same voxel. The original colors of the target barcodes blend with the other barcodes located nearby in the original physical space, making it difficult to accurately estimate the barcodes associated with cells in a given region of tissue. (Compare this figure with the high-resolution data shown in the middle panel of the top plot of Figure 7, where colors for individual axon targets can be clearly seen.)

**Figure 9:**
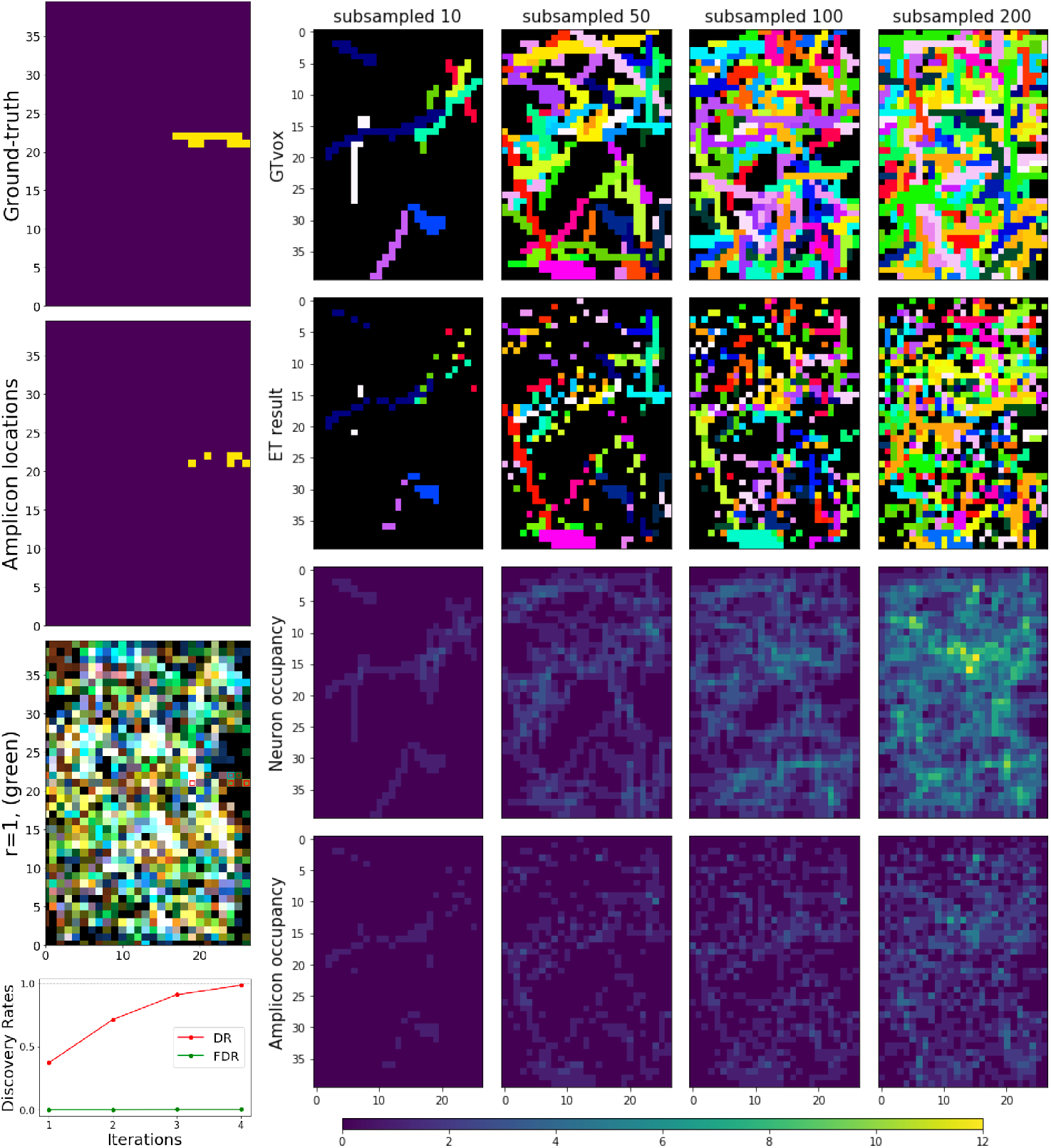
Limited optical resolution in low-resolution simulations yields signal mixing between neighboring neurons, but barcode discovery is still possible through an iterative process. Left, from the first to the third row: We show an example of one neuron. From the first to the third row: ground-truth morphology of the neuron; simulated amplicons; the image slice at the first imaging round. The red squares on the third row highlight true amplicon locations for a selected neuron, corresponding to the second row. Green fluorescence signal would be expected for the voxels containing amplicons with this neuron’s barcode in the first imaging round, but other colors are also present in these voxels due to contributions from other neurons. **Bottom Left:** The Discovery Rates (DR) result of the iterative barcode discovery process described in Section A.2.2. As we proceed through more iterations, the False Discovery Rate (FDR) remains essentially zero but many new correct barcodes are discovered. Four iterations were sufficient to find all the barcodes in this low-resolution simulations data. **Right:** Detailed look on the simulated low-resolution data. The neurons are highly overlapped in this data, so for visualization purposes we here show the results after subsampling the 412 neurons present in the data. Each column corresponds to a different number of subsampled neurons, and the number of randomly selected neurons are indicated in the title of each column. The first two rows show the ground-truth neuron segments and the evidence tensor (thresholded at 0.1) for the same neurons. As in Figure 7, each neuron segment is plotted with a distinct color. Note that the evidence tensor roughly captures the neuronal morphology shown in the first row. The last two rows show the number of overlapping ground-truth neuron segments and the number of amplicons for these neurons for each voxel. The high degree of overlapping can be seen.

#### 4.2.1 Barcode estimation

For this dataset, first we applied the same naïve barcode discovery approach that we used for the high-resolution simulations. This naïve method was able to find less than 40% of all the barcodes in the imagestack (assuming a maximum false discovery of 3 barcodes). Therefore, as in high-resolution high-density regime, we adopted an iterative approach. As shown in the bottomn plot of the left panel of Figure 9, four iterations of this approach were sufficient to find all the barcodes (412 in total) for this data, achieving 100% discovery rate with only 1 false positive. This result suggests that even in this more challenging low-resolution regime, an iterative approach can nonetheless recover the barcode library accurately.

#### 4.2.2 Amplicon estimation and morphological reconstruction

Neuronal tracing on this low-resolution simulation was performed using a similar approach to that taken on the high-resolution dataset. For this dataset, as shown in the top panel of Figure 10, the original evidence tensor already recovers the true amplicon locations quite accurately; a thresholded evidence tensor yields an adequate (though discontiguous) representation of the path of each neuron. Alphashape or CNN methods can be used to further improve this result, giving reasonable estimates for the contiguous space occupied by each neuron (see Figure 10). All together we see that we are still able to detect the correct morphology of the captured neurons, even for this low-resolution STIBE.

**Figure 10:**
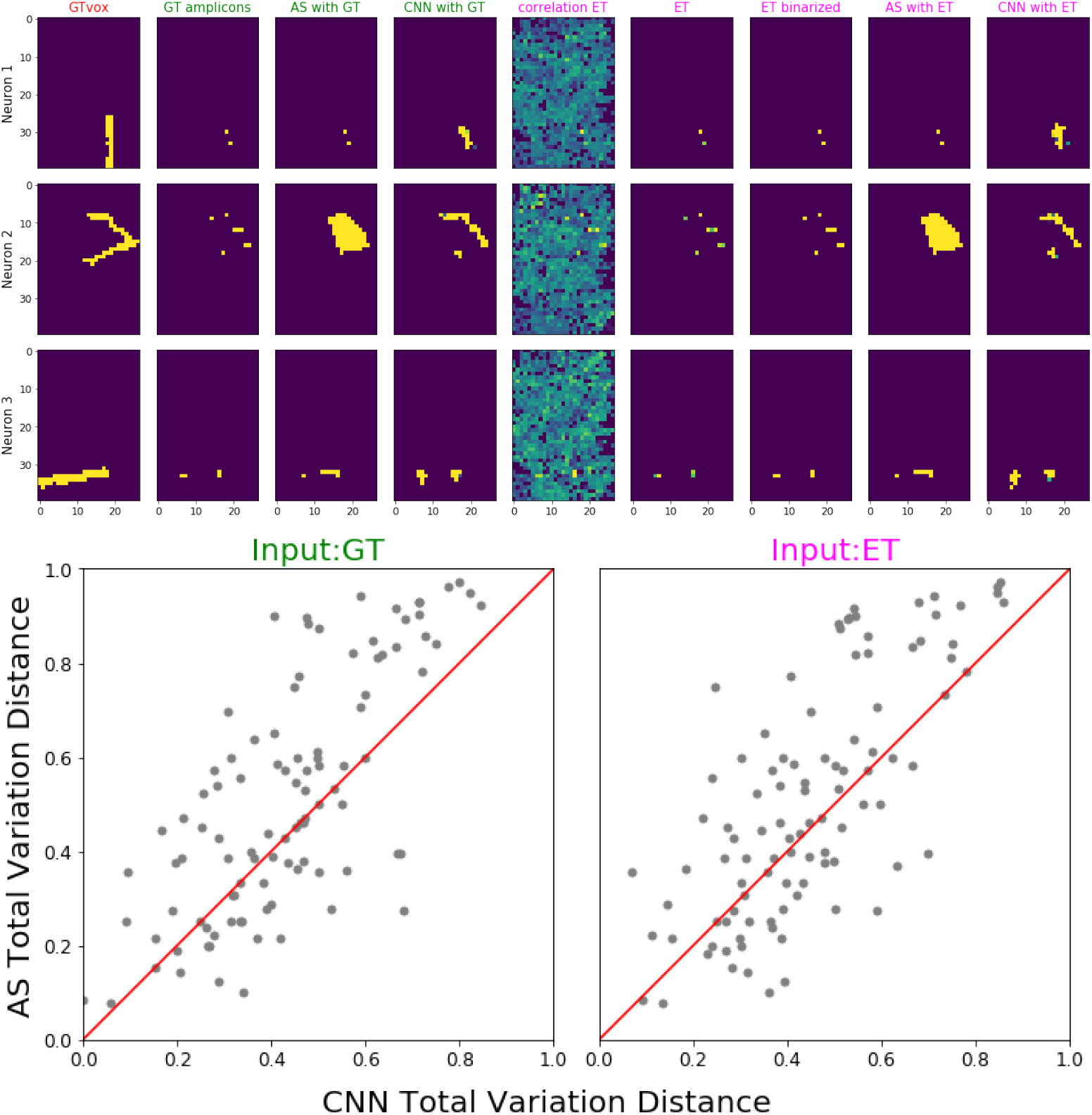
Morphology reconstruction results from CNN on low-resolution, sparse-expression simulations. Similar to Figure 8 for high-resolution simulation, this figure compares morphology reconstructions from low-resolution data using alphashape (AS) and CNN, with ground-truth (GT) amplicons and evidence tensor (ET) as input. Overall, we see that micron-precise morphological reconstructions are possible, even from low-resolution STIBEs. **Top:** Morphology reconstruction results, using either groundtruth amplicons or the evidence tensor as the input to alphashape or CNN. As in Figure 8 the colors of column titles indicate the input categories; from left to right the columns are: the ground-truth morphology, ground-truth amplicons, alphashape and CNN prediction provided ground-truth amplicons as input, evidence tensors using correlation methods, evidence tensors using amplicon density (described in Appendix A.3.1; note that this is much more accurate than the simple correlation-based method here), thresholded evidence tensor (using amplicon density thresholded at 0.1), and alphashape and CNN prediction given evidence tensor as input. For the alphashape input, the evidence tensor obtained from amplicon density was binarized with threshold 0.1. **Bottom**: Evaluating the morphology reconstruction using total variation distance loss (see Appendix A.3.2). On the left we show performance using ground-truth amplicon locations (GT, left) and on the right we show performance using the evidence tensor (amplicon density) derived from the imagestack (ET, right); conventions as in Figure 8. CNNs outperform alphashape modestly here.

## 5 Conclusion

In this work we developed detailed simulations to test the feasibility of using Spatial Transcriptomics-based Infinite-color Brainbow Experiments (STIBEs) for morphological reconstruction of many neurons simultaneously. We developed a novel blind-demixing algorithm, followed by a neural network based reconstruction approach, which together achieve high barcode detection rates and accurate morphological reconstructions, even in relatively low-resolution settings.

There may be room to improve further on the methods proposed here, on at least two fronts. In the first stage, we use an iterative approach to detect barcodes: we model the presence of barcodes that are already in our library, and search the residual images for new barcodes to add to our library. We found that the proposed linear programming approach (building on previous work described in [16]) was effective for decomposing the observed signal into the sum of these two parts (i.e., the signal that can be explained by barcodes we already know and the left over residual), but in future work it may be possible to improve further, using e.g. neural networks trained to decompose images using a simulator similar to the one used here [46].

Second, given an estimated barcode library, we would like improved methods for recovering the morphology corresponding to each barcode from the imagestack. One promising direction would be to use more sophisticated approaches for 3D neuronal recovery, adapting architectures and loss functions that have proven useful in the electron microscopy image processing literature [47,48]. We hope to explore these directions in future work.

## 6 Acknowledgements

We thank Xiaoyin Chen, Li Yuan, Tony Zador, and Abbas Rizvi for many helpful discussions. This work was supported by the National Institutes of Health (1U19NS107613), IARPA MICrONS (D16PC0003), and the Chan Zuckerberg Initiative (2018-183188).

## A Detailed methods

### A.1 Data simulation

The simulation process is illustrated in Figure 2. Specifically, we started with all the cells from a selected region of brain, densely-reconstructed from the MICrONS electron microscopy dataset (cf. MICrONS Explorer [14, 15, 43]). A (4 × 4 × 4) *μm* region is shown in Figure 2. These shapes were originally stored as triangular meshes; we converted them to binary 2D or 3D voxel tensors, using the voxels with pre-specified size under each simulation setting (the original MICrONS data was 3.58 × 3.58 nanometer resolution with 40 nanometer thick sections; in Figure 2 we show (100 × 100 × 100)-nanometer voxel size, as in the high-resolution simulations).

For each neuron we also created a unique ‘one-hot’ barcode, assuming *R* rounds of imaging with *C* channels. That is, for each neuron *j* we constructed a random binary matrix 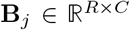 such that each row contains exactly one nonzero entry. Assuming *C* = 4 color channels, this enables labeling of 4^*R*^ unique neurons; we used *R* = 17 in all our simulations. Three examples of simulated barcodes can be seen in the top right panel in Figure 2.

Our assumption that each cell carries a unique barcode merits discussion, on two points. First, two neurons might be labeled by the same barcode, or very similar barcodes. This potential issue was discussed in a recent review [44], where it is shown that the fraction of uniquely labeled cells approximates to 1 when the number of barcodes is sufficiently large in relative to the number of cells that are labeled. Since in our simulation study, the number of possible barcodes (4^17^) is much larger than the number of cells (which is in the magnitude of 10^2^), we concluded that it was safe to assume that every neuron under study here is uniquely labeled. Indeed, after the barcodes are uniformly sampled, we found that the minimum Hamming distance between any pair of barcodes is at least 8, which is enough to easily distinguish one from the others in the imagestack. Second, a single neuron might be labeled by two distinct barcodes (‘double-infection’), though previous work indicates this is rare [9]. For simplicity we here ignored this possibility; if double-infection were a significant problem, it would be necessary to implement an algorithm to match pseudocolors together if they trace out similar paths. We leave this for future work.

To construct the imagestack itself, we input the transcript locations and codebook into Algorithm 1. This algorithm attempts to replicate the physical processes by which transcripts in tissue become fluorescent signals which are captured by a camera. It includes parameters to control many features of this process, including the variability in overall signal strength of each transcript, variability in per-round signal strength, a point spread function (PSF), and random per-voxel ‘speckle noise.’ We set ‘signal_range’ to be (10, 15) (randomly scaling the overall signal strength of each amplicon between 10 and 15). We set ‘per_frame_signal_range’ to be (0.8, 1) (randomly rescaling the signal in each frame independently for each amplicon between 0.8 and 1.0). We based these parameters on manual investigation of recent experimental data [6]. We used several different choices for ‘blursz,’ depending on the dataset simulation details (see the following subsections).

#### Algorithm 1: create_imagestack

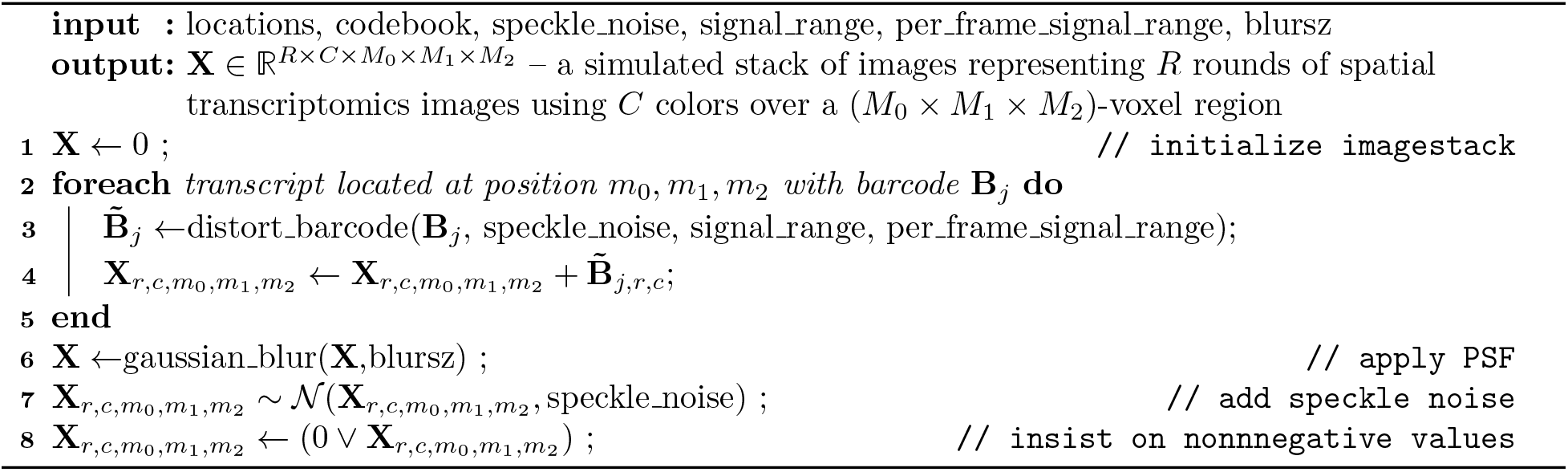

#### Algorithm 2: distort_barcode

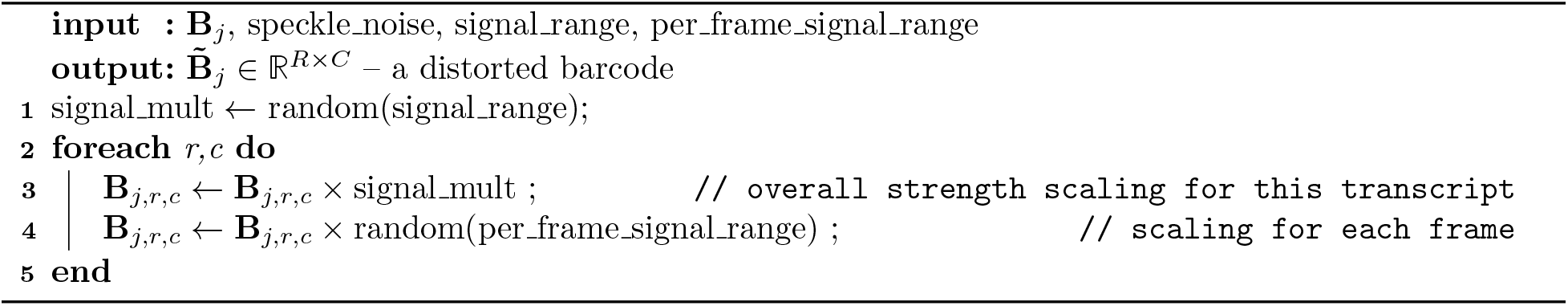

**Table 1:** Notation

|  | Description | Dimensions | Support |
| --- | --- | --- | --- |
| $M_0, M_1, M_2$ | number of voxels in each dimension (product is $M$) | scalar | $\mathbb{N}$ |
| $C$ | number of types of probes/wavelengths | scalar | $\mathbb{N}$ |
| $R$ | number of imaging rounds | scalar | $\mathbb{N}$ |
| $J$ | number of barcodes used to label neurons | scalar | $\mathbb{N}$ |
| $\mathbf{X}$ | simulated imagestack | $R \times C \times M$ | $\mathbb{R}^+$ |
| $\mathbf{B}$ | binary codebook matrix | $J \times R \times C$ | $\{0, 1\}$ |
| $\tilde{\mathbf{B}}$ | distorted binary codebook matrix | $J \times R \times C$ | $\{0, 1\}$ |
| speckle_noise | noise added to the simulated imagestack | scalar | $\mathbb{R}$ |
| signal_range | range of the intensity for each frame, each amplicon | vector | $\mathbb{R}^+$ |
| per_frame_signal_range | fluctuations on the signal strength for each frame | vector | $[0, 1]$ |
| blursz | size of PSF in units of voxels | vector | $\mathbb{N}$ |
| round_satisfaction_threshold | threshold for a channel to be considered to be active | scalar | $[1, \infty)$ |

#### A.1.1 High-resolution, dense-expression simulation

For the high-resolution simulation setting, for every neuron in a selected region of the brain (one (14.6 × 14.6 × 19.7) *μm* field of view (FOV) from the MICrONS dataset), we first voxelized the mesh objects using (100, 100, 100) *nm* voxels. Then we simulated the locations of amplicons (barcoded RNA transcripts) inside that voxelized neuron. These locations were sampled from a homogeneous Poisson process with rate λ. This rate constant has units of ‘density per cubic microns’; it indicates the average number of amplicons one would expect to find in a given cubic micron of neuronal tissue. See Figure 3 for examples of several imagestacks simulated with different densities λ. If λ is too low, the barcode signal will be too sparse to reconstruct or even detect cells; if λ is too high, the signal could become overwhelmingly mixed during imaging, making it difficult to identify the boundaries between multiple neurons in close proximity. See Figure 4 for an illustration of this density issue, even with a medium amplicon density.

Once we have determined the numbers and the locations of these amplicons, we applied a PSF to the images to mimic the image obtained from light microscopy; we used a Gaussian kernel with standard deviation of size (200, 200, 200) *nm*, blurring the signal by roughly 2 voxels in all directions.

#### A.1.2 Low-resolution, sparse-expression simulation

For the low-resolution simulation, the simulation process was similar to the high-resolution case described above, with some modifications. The first difference is that instead of studying one FOV, we started from a larger region of the brain (of size (196, 129, 40) *μm*, which is the full region used in [43]). Second, the voxel size used to convert the original mesh data was (5 × 5 × 80) *μm*, and the PSF was not added for this simulation since the optical resolution is so low in this setting. The number of amplicons in a neuron is considered to be proportional to its axon length (amplicons per micron length), instead of volume of the axons (amplicons per cubic micron): we used 0.08 amplicons per *μm* as the density of the simulated amplicons, matching [5]. Also, only the axonal segments are considered, and we used a simple criterion to remove dendrites (described in A.3.5). This is because in this setting we are simulating the tracing of long-range neuronal projections, and therefore restricted our focus to only the axonal parts of the neuronal segment rather than dendritic parts, even though the original EM data included all the neurons within the selected region. Lastly, only a small proportion of neurons were modeled to be infected by the virus (i.e., labeled by the barcodes). We randomly selected only 1% of all the neurons from the EM images (a more realistic assumption based on current cellular barcoding technologies [5]). An example of the resulting simulated data is shown in the left panel of Figure 9.

### A.2 Barcode estimation

Identifying the barcodes present in a given region of tissue is straightforward in settings with high optical resolution and low labeling density, because we can take advantage of the one-hot nature of the barcodes: each barcode is designed so that in each round exactly one channel is illuminated. On the other hand, for low-resolution and/or densely-labeled datasets, a more sophisticated iterative approach is necessary. We outline the main ideas below.

#### A.2.1 Naïve barcode library estimation

If a voxel contains signal dominated by only one barcode, exactly one channel should be brightest in each imaging round. This suggests the following procedure: scan for voxels that are ‘nearly one-hot’ — i.e., in each round there is a single channel which is much brighter than all others — and use the bright channel in each round to estimate the barcode for this voxel.

To make this idea practical, there are several difficulties that must be overcome. First, we need to find a way to look at the fluorescence data and decide when a voxel is ‘nearly one-hot.’ For this it is necessary to pick a threshold value. Second, we need a way to integrate information across the entire tissue. We can collect barcode candidates from a modestly large region of tissue, but each barcode candidate may be corrupted by noise from the data, yielding many almost-identical barcodes that are all slightly incorrect. To get the best possible barcode estimates, we need a way to merge similar barcodes together.

We adopt the following strategy. At each voxel location *m* and for each round *r* we identified the brightest channel, 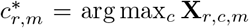. We also identify whether this brightest channel is significantly above the other channels. Using this information we produce an initial estimate of a barcode that might be present in this voxel. These initial barcodes are defined by thresholding:

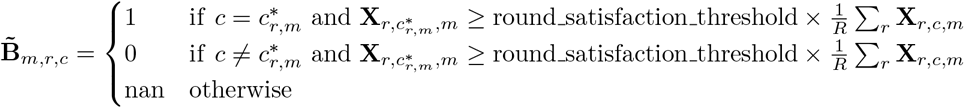

Here, a ‘nan’ indicates that we are uncertain about the barcode signal for this location at this round; i.e., the signal from the brightest channel 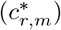 is not high enough for us to confidently determine this is the correct channel. ‘Round_satisfaction_threshold’ is set to be 1 (i.e., the bright channel must be higher than the average of all the rounds) in the results presented here. We compared the signal strength in these bright channels with the total signal strength, according to the formula

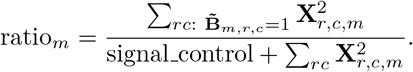

Here ‘signal_control’ is a parameter that controls our sensitivity to voxels with low overall strength; unless total signal strength in **X** at voxel *m* is significant, this term will cause the overall ratio to become small. For each voxel location *m*, if ratio_*m*_ is higher than a threshold, we conclude there is a valid amplicon at this voxel location.

We construct an initial codebook by concatenating the codebooks associated with all the voxels where the ratio discussed above exceeds a threshold. We then attempt to remove duplicates from this codebook. We first merge any barcodes that are exactly identical. We then use a greedy algorithm to merge any barcodes which are within a given Hamming distance (we chose a minimal distance of 3 here, as we found that amplicon locations could still be accurately estimated with this level of corruption). Barcodes are merged according to the formula

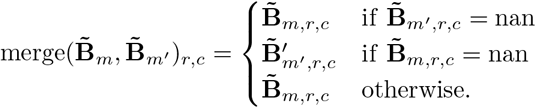

Note that this approach uses the first barcode’s information if there is disagreement: the order in which barcodes are merged could make a difference. So far this has not introduced any significant problems, but there are alternatives that could be pursued in the future (e.g., ordering so that the brightest barcodes come first).

Henceforth we will use **B** to denote the final codebook yielded by the approach above, with **B**_*j, r, c*_ denoting the value for barcode *j* in round *r* and channel *c*. We will let *J* denote the total number of barcodes identified (after deduplication).

#### A.2.2 Iterative barcode discovery and amplicon location estimation

In low-resolution or densely-labeled settings, the naïve approach outlined above was not sufficient. There were many barcodes which did not appear alone in any voxel, making it essentially impossible to identify them using the method above. In this case, we found the following iterative procedure effective.

We first identify some barcodes in the imagestack, using the naïve approach given above. We then estimate signal arising from those barcodes, and attempt to remove it from the imagestack. We then apply the naïve approach again to the residual imagestack to find additional barcodes. Next, we return to the original imagestack, and attempt once again to estimate all signal arising from any barcode that we have discovered. This procedure, formalized in Algorithm 3, can be repeated until no new barcodes are discovered (or for a prespecified number of iterations).

##### Algorithm 3: iterative_barcode_and_amplicon_estimation

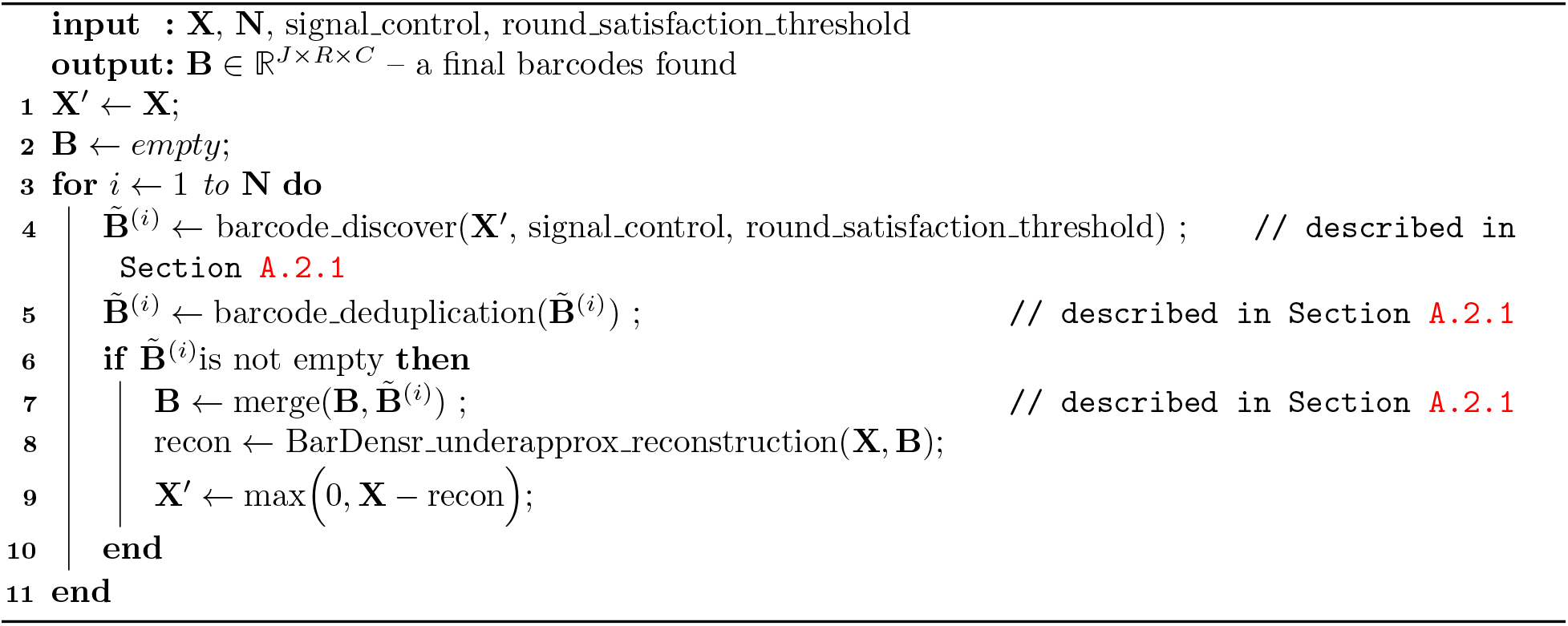

To enact this algorithm, we need a way to estimate all signal in an imagestack arising from a particular set of barcodes, *without* full knowledge of the true barcode library. For this task we build on the BarDensr model [16], which demixes and deconvolves spatial transcriptomic imagestacks by modeling the physical process that gives rise to the observed fluorescences.

We assume the following non-negative matrix factorization model for the observed data, **X**:

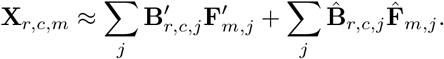

Here 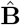 represents the set of barcodes which have already been discovered, 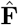 indicates a non-negative fluorescent intensity for each known barcode at each voxel, **B**′ represents the set of barcodes which are yet undiscovered, and **F**′ indicates a non-negative fluorescent intensity for each unknown barcode at each voxel. Note that we do not expect the equation above to hold exactly, due to various forms of noise, but we hope that it will be approximately correct. This equation can be seen as a version of BarDensr model, generalized to the case that some barcodes are unknown.

Under this model, the signal from the known barcodes corresponds to 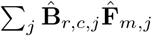. To estimate this signal it is thus sufficient to estimate 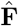. This problem is difficult because of the the unknown barcodes contributing signal to the observed imagestack **X**. Fortunately, we can make use of the non-negativity constraints of the model to help. From these constraints, it follows that

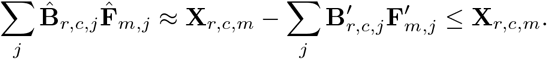

We call this the ‘under-approximation’ constraint, because it ensures that the estimated signal arising from the known barcodes is less than the observed fluoresence.

Now we need to choose an appropriate loss function. One approach would be to find 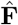 by minimizing the mean squared error between **X**_*r, c, m*_ and 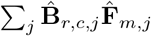; this would be similar to the approach taken in [16], where a quadratic objective together with a simple non-negative constraint could be efficiently optimized using standard techniques from the non-negative matrix factorization literature. However, adding the under-approximation constraint leads to more generic kinds of quadratic programming problems which are expensive to solve when the number of voxels is large. By slightly modifying this objective, we can obtain a fairly easy linear programming problem: instead of minimizing the mean squared error, we maximize the dot-product between the data and the estimated signal from the known barcodes. Thus we want to account for as much of the observed fluorescence as possible without overstepping the constraints. (Mathematically, we have replaced a quadratic penalty on the reconstruction with a linear penalty and a set of linear constraints that bound the magnitude of the reconstruction at each voxel.)

Putting the constraints together with the loss function, we arrive at the following approach for estimating the signal from the known barcodes. We first solve the following linear programming problem.

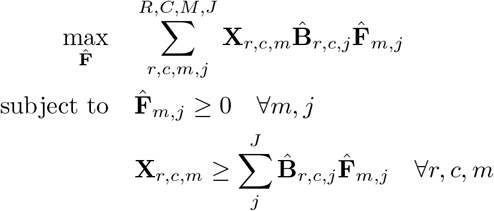

We then take 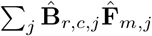 as our estimate of the signal arising from the known barcodes. We call this problem the ‘BarDensr_underapprox’ problem. This new problem enforces non-negativity in two ways: it enforces that 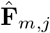, is non-negative but it also enforces that 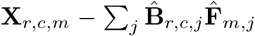 is non-negative. This approach is fairly fast, especially with parallel hardware such as GPUs, because the main computational effort comprises linear programming problems which can all be solved independently. We found we could make it faster still by assuming that only a small subset of the fluorescence densities were nonzero at each voxel. In particular, at each voxel we computed the correlation between that voxel and each barcode; we then assumed that only nine barcodes with the largest correlations carried nonzero fluorescent densities. This approximation yielded essentially no difference in the final results on any data that we tried it on. In practice, we found we could solve the ‘BarDensr_underapprox’ problem about six times faster than the original quadratic BarDensr optimization problem (16 seconds versus 96 seconds for a (30,30,30)-voxel patch using an Nvidia T4 GPU).

We found this constrained optimization approach to be effective even when our estimate of the barcode library represents a small fraction of the true set of barcodes which give rise to the imagestack. We also investigated a variant of this method which does not use the under-approximation constraint; this variant was unsuccessful. This finding adds to a growing body of work in the non-negative matrix factorization literature that shows that imposing under-approximation leads to robustness in the face of unknown contaminating signals [49, 50].

### A.3 Amplicon estimation and morphological reconstruction

Having estimated the barcode library, we must next estimate the amplicon locations for each barcode. Finally, we must use those amplicon locations to reconstruct the neuronal morphology, by ‘connecting the dots’ between each estimated amplicon ‘dot.’

#### A.3.1 Constructing an ‘evidence tensor’ to match barcodes to voxels

To begin, it is convenient to compute, for each voxel and each barcode, some estimate of our confidence that this barcode appeared in this voxel. We call an object of this kind an ‘evidence tensor.’ For the two simulations presented in this paper, we use two slightly different approaches to obtain such a tensor.

High-resolution, dense-expression simulation. We used a correlation method to compute the evidence tensor for each voxel and each barcode found:

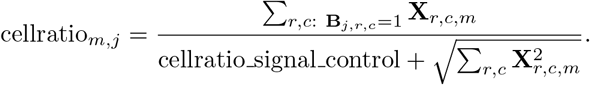

These ratios give a rough indication for whether a cell with barcode *j* might be present at voxel *m*. Where they are higher it suggests such a cell may be present. An example of the evidence tensor on a 2D plane is shown on the top panel of Figure 7, where a good agreement between the correct neuronal shape and the structure of the evidence tensor is seen.

Note that although this metric is relatively fast to compute, to directly apply it to the entire array of high-resolution data (size of (146, 146, 197) voxels, which gives *M* = 4,199, 252 voxels) for all neurons found (which gives *J* = 975 if all the barcodes are found) was not feasible because of memory constraints. Therefore, we used a simple tiling method and computed the evidence tensor for every discovered barcode within multiple small tiles with size of (30,30,30) voxels, with 10 voxels overlapping at each coordinate. We can further scale up the computation to even larger FOVs by additionally including the knowledge that there is only a small portion of the discovered barcodes present within each small tile.

Low-resolution, sparse-expression simulation. The simple correlation method above is insufficient analyzing low-resolution data; in this dataset the typical voxel has contributions from many barcodes, and the correlation method does not appear to be effective in this situation. Instead, we use ‘amplicon density’ as the evidence tensor, as described in [16]. Briefly, using BarDensr, the original imagestack **X** is demixed into two components - the amplicon density **F** and the accumulated barcode signal **G** (which is the scaled version of **B** mentioned in A.2.2):

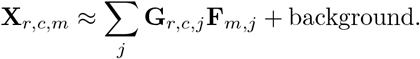

Similar to cellratio_*m, j*_, the amplicon density **F**_*m, j*_ at voxel *m* and barcode *j* indicates whether this barcode is present at this location.

#### A.3.2 Using the evidence tensor to reconstruct morphology

The evidence tensor constructed above is at best a noisy indicator of the neuron’s morphology. On the one hand, there may be voxels inside the cell without any transcripts. On the other hand, there may be voxels influenced by several cells (due to PSF blurring, voxelization artifacts, errors in barcode or amplicon estimation, etc.). To use the evidence tensor to best advantage in estimating the final morphology, we consider two approaches.

First, we considered alphashape [45], which is a widely used approach to estimate solids from pointclouds. The method takes one parameter, *α*, used to balance over- and under-coverage of the resulting estimate. Visual inspection was used to select this parameter to obtain the best possible result. For high-resolution simulations we used *α* = 0.1, and for low-resolution simulations we used *α* = 0.05.

The second approach involves Convolutional Neural Networks (CNNs). We used our simulation algorithms to generate an endless supply of imagestacks in which the ground-truth neuron morphology was known. We then trained a neural network to reconstruct the true morphology of a single neuron from the evidence tensor for the corresponding barcode. We used a four-layered CNN with residual blocks structure [51]. The network was trained on ‘total variation distance’ as the target function:

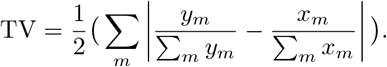

Here, *x* and *y* are either integer or float vectors. In our case, *y_m_* ∈ {0, 1} is the ground-truth label at voxel *m* and *x_m_* ∈ [0, 1] is the predicted label for *m*. To train the network while ensuring the relevant rotational symmetries, we used data augmentation. More details can be found in Appendix A.3.4.

#### A.3.3 Evaluating performance

We use two different criteria to measure the performance of our methods. The first is the discovery rate, which quantifies the proportion of the neuron barcodes that we can identify. The second criterion was total variation distance, described above, a quantitative way to represent the accuracy of morphological reconstruction.

#### A.3.4 Tiling and Data Augmentation for Convolutional Neural Net training

To train neural networks to reconstruct morphology from evidence tensors in the high-resolution setting, we used the procedure discussed above to generate images of size (60, 60, 60). We then augmented the data by taking six transformations of each tile: flipping over each axis, and swapping each pair of axes. All of these transformed versions of the data were fed as training data to the network at the same time, which helped ensure the network learned a function which was invariant to these kinds of transformations.

For network training in the low-resolution setting, we generated images of size (40, 27), and augmented this data with three transformations: flipping over each axis and swapping the two axes.

#### A.3.5 Euclidean Distance Transform (EDT)

For low-resolution simulation, we used Euclidean Distance Transforms (EDT) to calculate the largest sphere that could be contained inside a neuronal segment, which is then used to determine whether a given segment was dendrite or axon; if a segment can contain a sphere with radius of 5 voxels (25 microns), we categorized it to be dendrite.

The Euclidean Distance Transform is defined as follows. One begins with a 2D or 3D image *f*, and let *S* be the set of voxels in the image *f* whose values are 1. Following [52], the distance map of the input image *f* is defined as

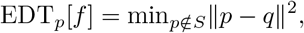

where ∥*p* – *q*∥^2^ is Euclidean distance between voxel locations *p* and *q*. The computational cost for computing this transform scales linearly with the number of voxels in the image [52].

